# Deficit in parietal memory network underlies auditory hallucination: a longitudinal study

**DOI:** 10.1101/204008

**Authors:** Qian Guo, Yang Hu, Botao Zeng, Yingying Tang, Tianhong Zhang, Jinhong Wang, Georg Northoff, Chunbo Li, Donald Goff, Jijun Wang, Zhi Yang

## Abstract

Auditory hallucination is a prominent and common symptom in schizophrenia. Previous neuroimaging studies have yielded mixed results of its brain network deficits. We proposed a novel hypothesis that parietal memory network, centered at the precuneus, plays a critical role in auditory hallucination. This network is adjacent and partially overlaps with the default mode network, and has been associated with brain function of familiarity labelling in memory processing. Using a longitudinal design and a large cohort of first-episode, drug-naïve schizophrenia patients, we examined this hypothesis and further investigated whether the functional connectivity patterns of the parietal memory network can serve as a neuroimaging marker for auditory hallucination and help to predict future treatment effects. Resting-state scans from 59 first-episode drug-naïve schizophrenic patients (27 with and 32 without hallucination) and 53 healthy control subjects were acquired at the baseline test, and 56 of them were scanned again after two months. Functional connectivity strength within the parietal memory network and between this network and memory hubs was across the three groups at baseline and follow-up scans. Results showed that decreased functional connectivity strength within the parietal memory network was specific to the auditory hallucination group (p = 0.009, compare to the healthy subjects; p = 0.029, compare to the patients without hallucination), with the precuneus representing the largest group difference. The intra-network connectivity strength of the precuneus negatively correlated with the severity of hallucination at the baseline scan (r = −0.437, p = 0.029), and it was significantly increased after two-month medication (p = 0.039). Logistic regression analysis and crossvalidation test demonstrated that the functional connectivity strength of the precuneus and precuneus-hippocampus connectivity could differentiate patients with or without auditory hallucination with a sensitivity of 0.750 and a specificity of 0.708. Moreover, crossvalidation test showed that these imaging features at the baseline scan well predicted the extents of positive symptom improvement in the hallucination group after the two-month medication (R^2^ = 0.433, p = 0.022). Our results provide evidence for a critical role of the parietal memory network underlying auditory hallucination, and further propose a novel neuroimaging marker for identifying patients, accessing severity, and prognosis of treatment effect for auditory hallucination.

**Abbreviations:**

AH: auditory hallucination
AHRS: Auditory Hallucination Rating Scale
AUC: Area-under-curve
BPRS: Brief Psychiatric Rating Scale
DMN: default mode network
DSM-IV: Diagnostic and Statistical Manual of Mental Disorders, Fourth Edition
DUP: duration of untreated psychosis
FCS: functional connectivity strength
HC: healthy control
IC: independent component
PMN: parietal memory network
NMDA: N-methyl-D-aspartate
RSN: resting state network
rs-fMRI: resting-state functional MRI
rTMS: repetitive transcranial magnetic stimulation
SANS: Expanded Version and the Scale for Assessment of Negative Symptoms

## Introduction

Approximately 70% of schizophrenia patients experience auditory hallucinations (AHs) through their lifetime course of illness (Sartorius *et al.*, 1974; Waters *et al.*, 2012). AH, defined as perception of sound in the absence of an external stimulus, can severely affect daily life functioning of schizophrenic patients. Insofar as numbers of neuroimaging studies have examined brain activation between healthy people and patients with or without AH, neural circuits underlying AH remain inconclusive.

As AH usually intrude into patients’ consciousness with no omen, there is a logical speculation that an atypical resting state may give rise to a hallucinatory experience. Resting state networks (RSNs), which reflect the spontaneous neural activities and patterns of connectivity between brain regions, currently draw most attentions in studies on phenomenon of AH. Among the notable networks at rest, default mode network (DMN), is regarded to be potentially relevant to the onset of psychopathology in general, and psychosis in particular (Alderson-Day *et al.*, 2015). However, literature is mixed to elucidate its possible contribution to AH. Jardri’s group compared state of hallucination (auditory hallucination, visual hallucination, or both) with “real rest” periods in 20 adolescents with brief psychotic disorder, and reported hallucinations were associated with disengagement and instability of the DMN, specific to modality (Jardri *et al.*, 2013). Other studies observed inconsistent DMN alterations in samples with auditory hallucination. Wolf discovered no difference in DMN function or correlation with symptoms, but observed alterations in precuneus and posterior cingulate, which belonging to an executive control network and a left frontoparietal network according to their analysis (Wolf *et al.*, 2011). In contrast, van Lutterveld *et al.* (2014) found increased connectivity in temporal cortices and the posterior cingulate/precuneus, representing posterior regions of the DMN, in a sample of nonclinical AH participants.

Such mixed results suggest that a possibility that subsystem of DMN or adjunct regions to DMN are associated with AH, and the above-mentioned cingulate/precuneus is a candidate given its critical role in memory processing (Cabeza *et al.*, 2008) and in coordinating multiple networks (Utevsky *et al.*, 2014). Indeed, a precuneus-centered network has been proposed by both task activation (McDermott *et al.*, 2009; Nelson *et al.*, 2010; Kim, 2013) and functional connectivity studies (Doucet *et al.*, 2011; Power *et al.*, 2011; Yeo *et al.*, 2011), which consistently reported synchronized intrinsic activity in precuneus, middle posterior cingulate (posterior part) and dorsal angular gyrus. These regions have been recently proposed to comprise the parietal memory network (PMN) that is strongly related to learning and memory and exhibits functional independence from the DMN in our previous work (Yang *et al.*, 2014) and other studies (Yeo *et al.*, 2011; Gilmore *et al.*, 2015).

Evidence from multiple modalities implicated that the PMN plays important role in memory encoding and retrieval, and is related to “familiarity labeling” in memory and learning (Nelson *et al.*, 2013). This function connects to one of the most influential theories of AH, that an unstable memory may trigger spontaneous and erroneous language production to cause AH (Hoffman, 1986). Furthermore, PMN may interfere with memory functions by interacting with classical memory regions such as hippocampus (Nelson *et al.*, 2013). Previous studies have demonstrated that associative memory performance can be persistently enhanced by stimulating dorsal parietal regions (where precuneus locates) using transcranial magnetic stimulation (rTMS), suggesting that the PMN is involved in memory and that PMN is functionally connected to hippocampus (Wang *et al.*, 2014). Hence, the precuneus-hippocampus functional connectivity may underlay AH generation.

In this study, we aim to investigate associations between PMN dysfunction and AH symptoms using resting-state fMRI. First-episode, drug-naïve schizophrenia patients with and without AH, as well as healthy participants, were compared at both baseline and follow-up scans after two-month medication. We examined whether AH symptoms are associated with deficient functional connectivity within the PMN and between the core region of PMN and the hippocampus. Further, we examined the power of these features to differentiate AH+ from general schizophrenic patients and to predict improvements of positive symptoms after treatment.

## Materials and Methods

### Participants

Fifty-three healthy controls (HC), 32 schizophrenic patients without auditory hallucinations (AH-), and 27 schizophrenic patients with auditory hallucinations (AH+) participated in this study. Two months after the baseline interview, 29 HC, 15 AH-, and 12 AH+ completed the follow-up interview and MRI scanning. All schizophrenia patients were recruited from Shanghai Mental Health Center, Shanghai, China. The healthy controls were recruited from local communities in Shanghai. Written informed consent was obtained from each participant or the participant’s guardian prior to data acquisition. This study was approved by the Local Research Ethics Committee.

**Table 1.** Baseline Demographic and Clinical Characteristics for First-Episode Schizophrenia and Healthy Controls

| Characteristics | First-Episode Schizophrenia |  | Healthy Controls<br>(SD) (N = 49) | Statistical Test (*P* value)<sup>b</sup> |
| --- | --- | --- | --- | --- |
|  | AH- Patients (SD)<br>(N = 28) | AH+ Patients (SD)<br>(N = 25) |  |  |
| Gender (Male/Female) | 10/18 | 12/13 | 26/23 | $\chi^2 = 2.16(0.34)$ |
| Age (years)<sup>a</sup> | 27.00(5.42) | 25.54(7.45) | 27.56(5.93) | F=0.82(0.44) |
| Education level(primary school/middle school/college) | 0/14/14 | 1/12/12 | 1/20/28 | Fisher's Exact Test=2.014(0.77) |
| Duration of untreated psychosis (weeks)<sup>a</sup> | 34.12(41.87) | 31.44(32.76) | — | t=0.76(0.45) |
| Auditory Hallucination Rating Scale<sup>a</sup> | — | 25.84(8.22) | — | NA |
| Brief Psychiatric Rating Scale<sup>a</sup> |  |  |  |  |
| Total score<sup>a</sup> | 46.82(8.70) | 46.68(8.29) | — | t=0.06(0.95) |
| Positive symptom score<sup>a</sup> | 15.32(4.33) | 16.36(3.84) | — | t=-0.92(0.36) |
| Assessment of Negative Symptoms<sup>a</sup> | 21.18(13.21) | 18.08(12.30) | — | t=0.88(0.38) |

<sup>a</sup> Data are presented as mean(SD).
<sup>b</sup> *p* values are in parentheses.

The inclusion criteria for the patient group were 1) consensus diagnoses of first-episode schizophrenia assigned by two psychiatrists, according to the Diagnostic and Statistical Manual of Mental Disorders, Fourth Edition (DSM-IV), on the basis of Structured Clinical Interview for DSM-IV; 2) medication-naïve; 3) education level higher than primary school and capability of finishing the tests; 4) age from 18 to 40. The exclusion criteria were 1) too agitated or aggressive to finish the assessments; 2) presence of another axis I psychiatric disorder; 3) rated 7 or higher in the Calgary Depression Scale for Schizophrenia (CDSS); 4) history of suicidal behavior; 5) history of antipsychotic medication; 6) history of substance abuse; 7) pregnancy; 8) history of serious physical diseases; 9) unsuitability for MRI scans, for instance, having metal implants. The HC group was matched with an SZ group for age, gender, and education level. None of the HCs had a positive family history for any psychiatric disorder. Participants were screened with Chinese version of the MINI, Version 5.0 (Sheehan *et al.*, 1998; Mei *et al.*, 2009) and excluded if they met criteria for any mental disorder according to the DSM-IV or had a history of serious physical diseases, pregnancy, taking any antipsychotic drugs, or substance abuse.

**Table 2.** Follow-up Clinical Characteristics for First-Episode Schizophrenia and Healthy Controls

| Characteristics | First-Episode Schizophrenia |  | Healthy Controls<br>(N = 25) | Statistical Test<sup>b</sup> |
| --- | --- | --- | --- | --- |
|  | AH- Patients<br>(N = 11) | AH+ Patients<br>(N = 10) |  |  |
| Gender (Male/Female) | 5/6 | 6/4 | 13/12 | $\chi^2=0.45(0.80)$ |
| Age (years)<sup>a</sup> | 24.27(4.71) | 25.80(5.39) | 28.84(5.86) | F=2.97(0.06) |
| *Clinical (First scan)* |  |  |  |  |
| Antipsychotic dose (mean CPZ equivalents, mg)<sup>a</sup> | 374.87(150.30) | 301.85(115.00) |  | t=1.20(0.25) |
| Auditory Hallucination Rating Scale<sup>a</sup> | – | 27.60(7.37) | – | NA |
| Brief Psychiatric Rating Scale<sup>a</sup> |  |  |  |  |
| Total score<sup>a</sup> | 50.09(8.11) | 48.30(7.48) | – | t=0.52(0.61) |
| Positive symptoms<sup>a</sup> | 15.27(2.01) | 18.00(3.23) | – | t=-2.35(0.03) |
| Assessment of Negative Symptoms<sup>a</sup> | 24.36(15.31) | 16.60(13.93) |  | t=1.21(0.24) |
| *Clinical (Second scan)* |  |  |  |  |
| Auditory Hallucination Rating Scale<sup>a</sup> | – | 8.50(11.36) | – | NA |
| Brief Psychiatric Rating Scale<sup>a</sup> |  |  |  |  |
| Total score<sup>a</sup> | 35.36(8.90) | 32.20(5.60) | – | t=0.98(0.34) |
| Positive symptoms score<sup>a</sup> | 8.82(4.36) | 8.50(3.72) | – | t=0.18(0.86) |
| Assessment of Negative Symptoms<sup>a</sup> | 19.73(12.28) | 13.90(9.69) |  | t=1.20(0.25) |

<sup>a</sup> Data are presented as mean(SD).<sup>b</sup> p values are in parentheses.

### Clinical measurements

A trained psychiatrist assessed the clinical symptoms of all patients. The baseline assessment was conducted before drug medication, and the follow-up assessment was conducted after 8 weeks of antipsychotic treatment. Each patient was clinically evaluated using the 24-item Brief Psychiatric Rating Scale (BPRS) Expanded Version and the Scale for Assessment of Negative Symptoms (SANS). For the schizophrenic patients with AH, the severity of AH was evaluated using a Chinese version of the auditory hallucination rating scale (AHRS) (translated by JW), which measures frequency, reality, loudness, number of voices, length, attention salience, and level of distress caused by the AH. These assessments were conducted at both the baseline and follow-up examinations. Duration of Untreated Psychosis (DUP) was acquired at the baseline.

### MRI data acquisition

All participants completed functional and structural MRI on a 3.0 T Siemens Verio MRI scanner (Siemens Medical Solutions, Erlangen, Germany) at Shanghai Mental Health Center. After 3-plane localizer, an anatomical scan was acquired with a T1-weighted magnetization prepared rapid gradient echo sequence (192 sagittal slices, echo time TR/TE/TI = 2300/2.96/900 ms, flip angle = 90°, FOV = 256 mm, matrix = 256×240, slice thickness/gap = 1.0/0.0 mm). An 8’30” resting-state fMRI was acquired subsequent to T1 image with an echo-planar imaging (EPI) sequence (45 axial slices, acquired from inferior to superior in an interleaved manner, TR/TE = 3000/30 ms, flip angle = 85°, FOV = 216 mm, matrix = 72×72, slice thickness/gap = 3.0/0.0 mm, 170 volumes). Subjects were instructed to close their eyes and remain awake during the MRI scan. Awakeness was confirmed in a brief interview after the scans.

### MRI preprocessing and quality control

Both structural and resting-state functional MRI (rs-fMRI) scans were pre-processed with the Connectome Computation System (Xu *et al.*, 2015). FreeSurfer (Dale *et al.*, 1999) was then used on structural data to extract brain and segment tissues into gray matter, white matter, and cerebrospinal fluid. All brain images were transformed into MNI152 standard space using ANTs (Avants *et al.*, 2011). As for rs-fMRI images, the following steps were applied: 1) discarding the first 15 volumes; 2) slice time correction; 3) head motion correction; 4) registration to corresponding structural images with rigid transformation and then converting into MNI space; 5) normalizing the data to a global mean intensity of 10000; 6) band-pass temporal filtering (0.01-0.1 Hz).

**Figure 1.**
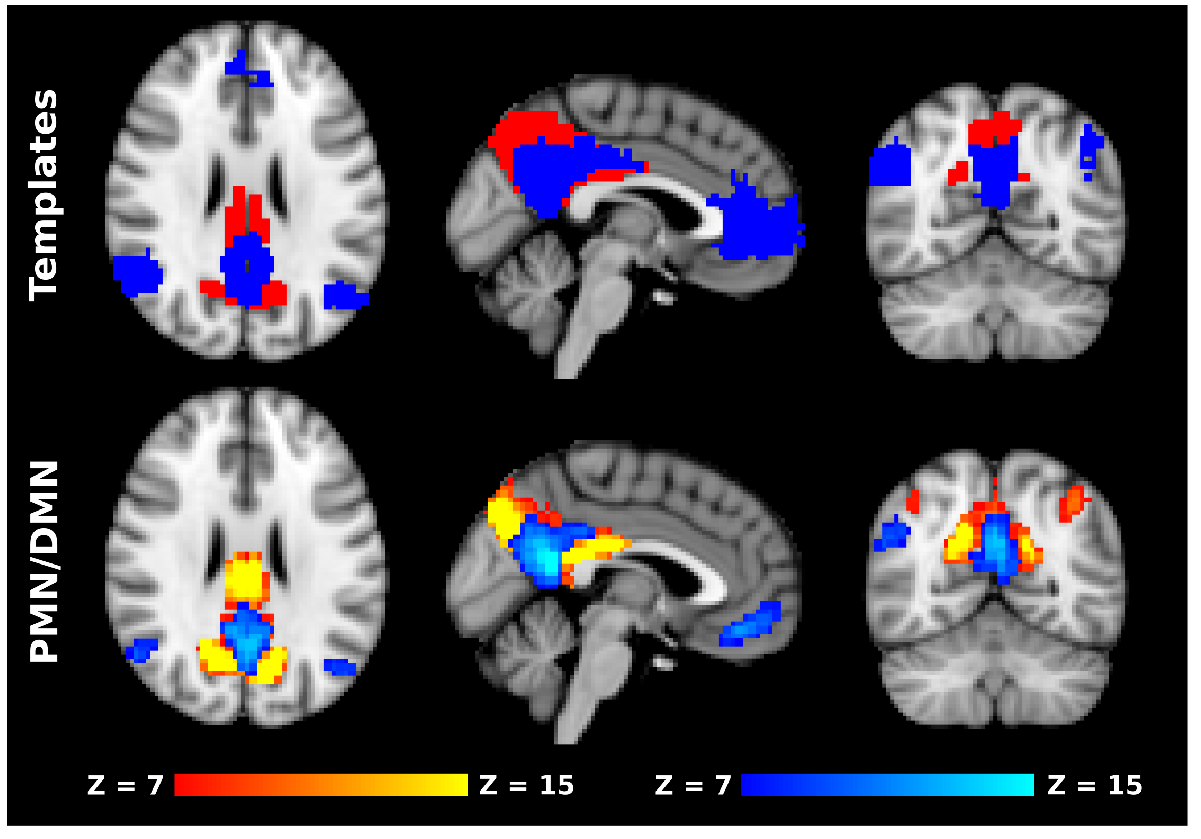
PMN and DMN are separated intrinsic networks. The upper row shows spatial distributions of PMN and DMN in intrinsic network templates obtained using clustering approach. The bottom row shows PMN and DMN identified in the present study. In both maps, cold color represents DMN and warm color represents PMN. There is a high similarity between the templates and the current findings.

The quality of brain extraction and spatial registration outputs was visually inspected. Images of 4 HC, 4 AH−, and 2 AH+ were excluded for further analysis due to poor brain extraction or spatial registration. The head motion in the rs-fMRI data were evaluated using mean frame-wise displacement (meanFD) (Power *et al.*, 2012), and all data had meanFD < 0.2 mm. Therefore, 49 HC, 28 AH−, and 25 AH+ entered the analysis of baseline scan, and 25 HC, 11 AH−, and 10 AH+ were included in the longitudinal analysis.

### Functional connectivity analysis

For the baseline scan, the pre-processed rs-fMRI images of all subjects were temporally concatenated and decomposed into a set of group independent components (ICs) using the MELODIC module of the FSL package (Beckmann *et al.*, 2005). The number of components was automatically estimated to be 51. ICs representing DMN and PMN were selected by matching against a pre-defined spatial template of resting-state networks by Yeo’s previous work (Yeo *et al.*, 2011). Dual-regression was applied to obtain individual IC maps and time courses for every subject (Beckmann *et al.*, 2009). In brief, for each subject, the spatial maps of the group-level ICs were used as regressors, and their contributions to the subject’s rs-fMRI data were estimated using a linear model. The contributions were depicted using time courses. In a same manner, these time courses were further used as regressors and their contributions to the same rs-fMRI dataset were estimated and represented as a set of spatial maps, each correspond to a group-level IC. These spatial maps represented resting-state functional networks (or image artifacts) in individual subjects.

### Statistical analyses

Statistical analyses of demographic and clinical data were examined using R (R Development Core Team, 2003). ANOVA models were used to compare continuous normally distributed variables among groups, and Chi-square tests were used for categorical variables.

For rs-fMRI data, we compared the ICs of interest across the three groups in two ways: 1) The IC maps for the DMN and PMN were thresholded at p = .500 using a Gaussian mixture model to reflect the core regions of the DMN and PMN. The mean of the voxel-wise weights within the core regions were compared across the three groups (Mingoia *et al.*, 2012). This metric reflects the overall functional connectivity strength (FCS) of a network because the weight of a voxel indicates its FCS with the core regions in the given IC. 2) To reveal foci with significant group differences, we further compared voxel-wise weights across the three groups using a nonparametric permutation test (5,000 permutations) for the DMN and PMN (Winkler *et al.*, 2014). The multiple comparison correction was conducted with the threshold-free cluster enhancement approach (TFCE) (Smith and Nichols, 2009). A region of interest was defined as a 6 mm-radius sphere centered at the identified foci with the maximal group difference. This ROI definition helps to alleviate the impact of spatial registration error. The mean weights of the ROI were further compared across groups and correlated with the AHRS scores in the AH+ group.

We further investigated the functional connectivity between the core ROI of PMN with bilateral hippocampi. The masks of bilateral hippocampi were determined using Harvard-Oxford subcortical structural atlas (fsl.fmrib.ox.ac.uk/fsl/fslwiki/Atlases). The mean signals of white matter/ventricles and six head motion parameters were regressed out from the ROI mean time series before computing functional connectivity. Correlation coefficients between time series were computed and transformed into Fisher’s Z scores to quantify functional connectivity. An ANOVA was performed to compare the functional connectivity measures across groups.

As for the follow-up dataset, we applied dual-regression again to project the rs-fMRI data onto the group-level IC maps that were obtained in the main analysis, yielding subject-specific ICs and their corresponding mixing time courses. Longitudinal analyses were conducted using repeated measure ANOVAs to examine changes of the mean weight of the PMN ROI and its interaction patterns with hippocampi. Changes of clinical scores were correlated to changes of functional connectivity. Age, sex, and education levels were included as covariates in all between-group comparisons in the above analyses.

**Figure 2.**
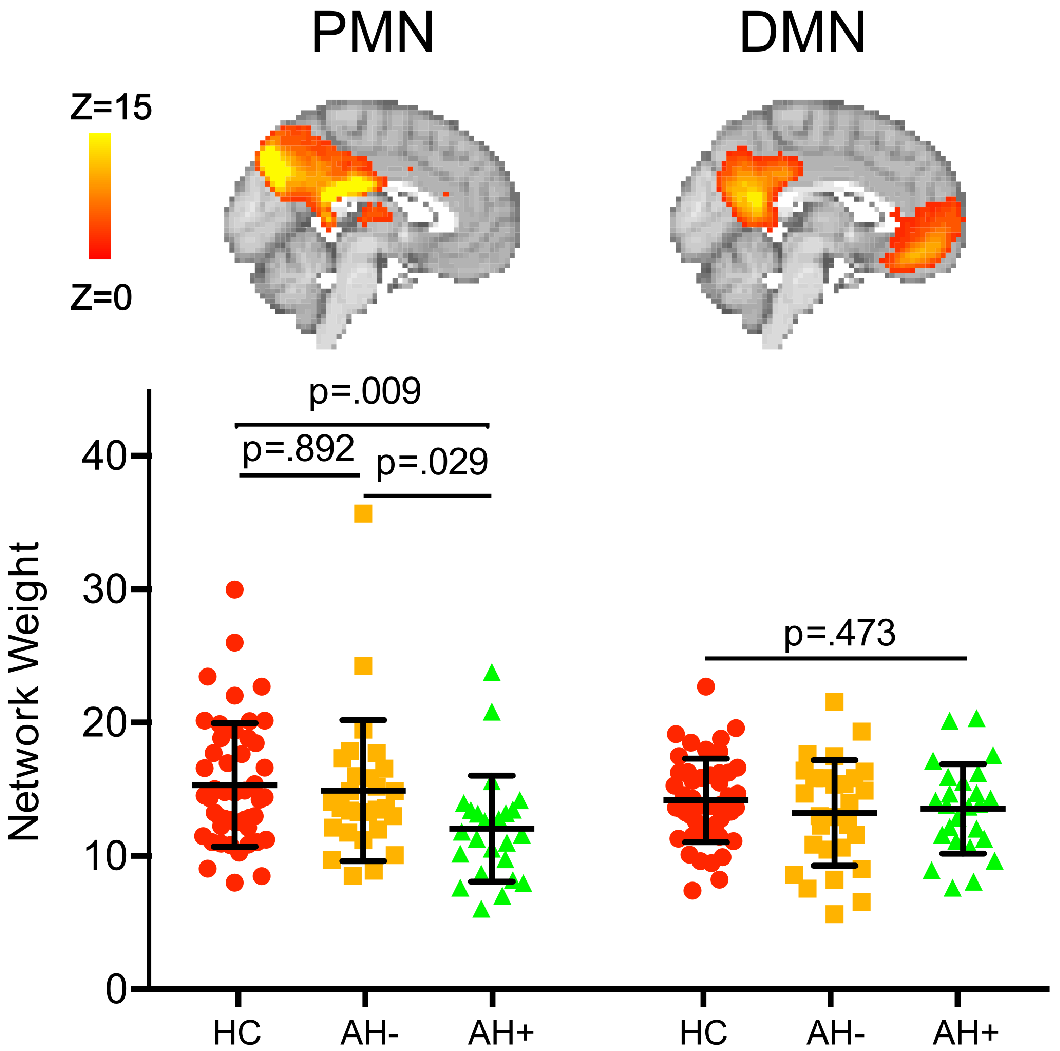
Network weights, representing overall functional connectivity strength, exhibits significant decrease in AH+ group in PMN. In DMN, there is no significant between group difference.

To further quantify the predictive power of the functional connectivity and clinical features to identify AH symptoms from general schizophrenia, we conducted a stepwise logistic regression between AH+ and AHȒ group. Area-under-curve (AUC) and leave-one-out crossvalidation accuracy were used to evaluate the predictive power of the regression model. We also examined a linear regression model to predict the improvement of positive symptom score of AH+ patients using the functional connectivity strength and clincal features.

## Results

### Demographic and clinical characteristics

At both baseline and follow-up, demographic data did not differ in age, gender, or education level among healthy participants, AH−, and AH+ patients (Tables 1 and 2). For clinical measures, AH− and AH+ showed no significant difference in BPRS total score, positive symptoms of BPRS, or SANS at the baseline. With two-month medication, AH+ showed significant decreases in AHRS (t = 6.449, p < 0.001), BPRS total score (t = 5.729, p < 0.001), and positive symptoms of BPRS (t = 5.003, p = 0.001), While AH− group exhibited significant longitudinal decreases in BPRS total score (t = 4.155, p = 0.002) and positive symptoms of BPRS (t = 5.212, p < 0.001). There was no significant difference in either the demographic or the clinical measures between the schizophrenic patients who were successfully followed and those who dropped out (Supplementary Table 1), supporting the representativeness of the follow-up dataset when comparing the AH+ to AH− groups.

### Relationship between PMN and AH symptom

The PMN and DMN component maps, identified by group-level ICA analysis, exhibited correlation coefficients of 0.47 and 0.44 respectively to resting-state network templates reported in Yeo’s work (Yeo *et al.*, 2011) (Figure 1). They were much higher than the second-best matching component maps (0.26 and 0.30, respectively). As presented in Figure 2, the three groups showed significant differences in the mean weight in the PMN (F = 4.263, p = .017), but not the DMN (F = .753, p = .473). Post-hoc analysis revealed significantly lower mean weight in the AH+ group, when compared to both HC (t = 2.650, p = .009) and AH− (t = 2.220, p = .029) groups. There was no significant difference between AH− and HC groups (t = -.136, p = .892).

### Functional connectivity strength of the core region in PMN reflects AH severity

Whole-network, voxel-wise analysis of variance revealed significant group difference in the right precuneus, a core region of the PMN, between the HC and AH+ groups (p = .009, 8 voxels, multiple comparison error corrected using the TFCE approach). No voxel survived in the comparison between the HC and AH− groups. To reduce the registration errors between baseline and follow-up MRI scans, a region of interest (ROI) was defined as a 6 mm-radius sphere centered at the peak position of above cluster (x = 6, y = -69, z = 27 in MNI space) for further statistical analysis (Figure 3A). When compared with the other two groups, AH+ exhibited significant decrease in regional mean weight (t = 3.553, p < .001 compared with the HC group; t = 2.430, p = .017 compared with the AH− group) as illustrated in Figure 3B. Moreover, the mean weight within this ROI showed a significant negative correlation with the AHRS scores (r = -0.437, p = .029, Figure 3C), linking alteration in the core region of the PMN to the severity of AH symptoms. A linear regression model including the mean weight of this ROI, age, gender, education level, and DUP of the patients explained 46.0% of the individual variability of AVHRS scores (R^2^ = 0.460, p = 0.057). The contributions of the mean weight of the precuneus ROI and age were both significant (p = 0.035 and p = 0.031, respectively).

**Figure 3.**
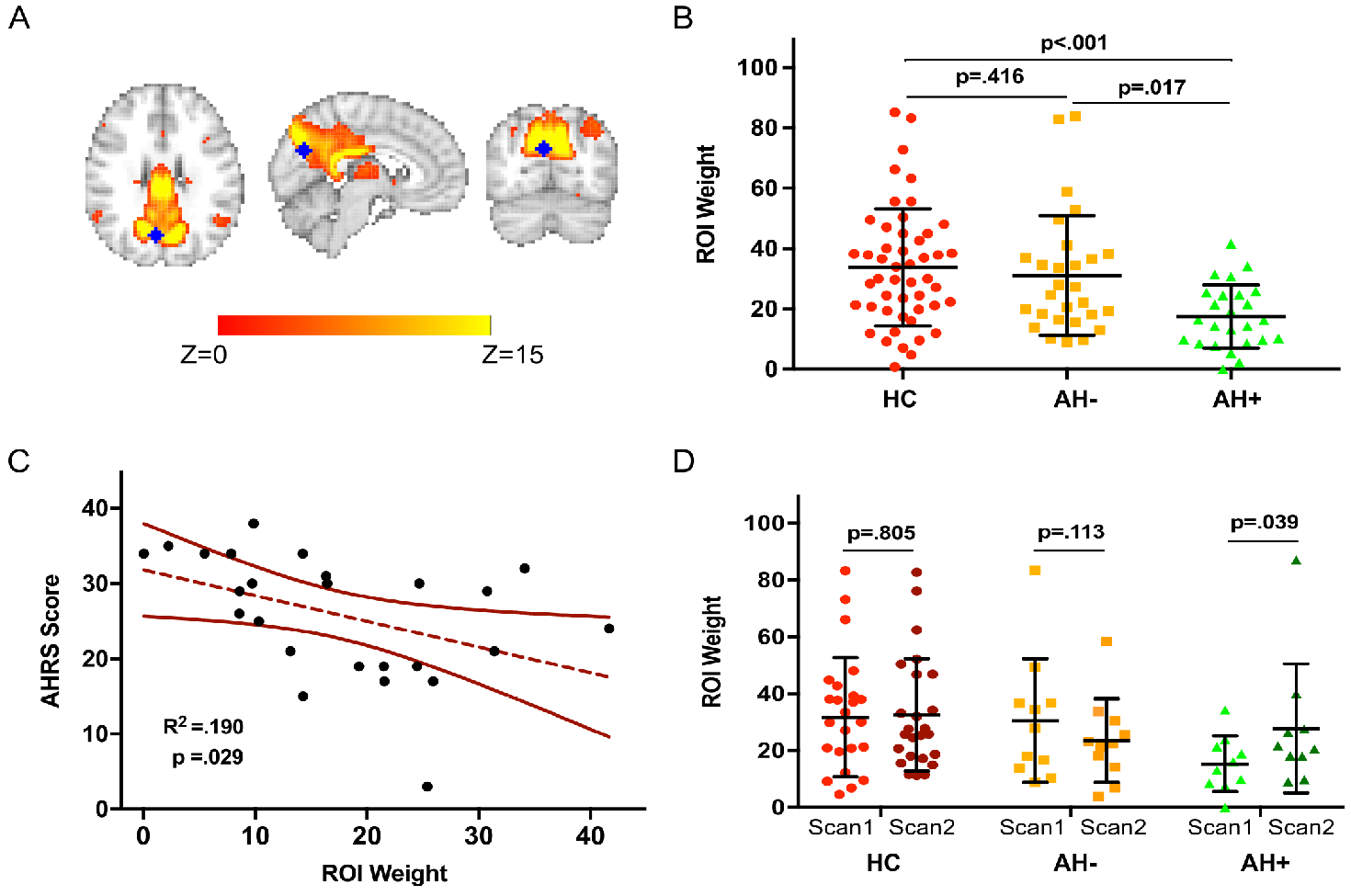
Overall functional connectivity strength in precuneus in PMN is relevant to AH symptoms, as reflected in multiple aspects. A. The location of the identified precuneus region; B. Functional connectivity strength in precuneus exhibits significant decrease only in AH+ C. Functional connectivity strength in precuneus negatively correlated with severity of AH symptoms; D. Functional connectivity strength in precuneus significantly increased after medication in AH+.

Longitudinal analysis (repeated measure ANOVA) revealed a significant group by time interaction in the mean weight of the precuneus ROI (F=4.365, p = .010). Simple effect analysis showed that only the AH+ group exhibited a significant increase in the mean weight of the ROI (t = 2.417, p = .039) after two-month medication (Figure 3D). This effect remained significant (t = 2.814, p = .023) when excluding an outliner in follow-up scan. There was no significant longitudinal change in either HC (t = .250, p = .805) or AH− groups (t = 1.739, p = .113). The changes of mean weight in the precuneus ROI did not correlate with the improvement of clinical scales in the AH+ group.

### Precuneus-hippocampus functional connectivity separate AH+ from AH−

A marginally significant group effect (F = 2.894, p = .060) was observed in FCS between the precuneus ROI and bilateral hippocampi. Since there was no significant laterality by group interaction (F = .075, p = .928) or laterality main effect (F = .062, p = .804), we averaged the FCS measures for the left and right hippocampi to yield a combined FCS between the precuneus ROI and hippocampus. AH+ showed significantly weaker FCS between the hippocampus and the precuneus ROI, compared to AH− (t = −2.577, p = .013), whereas the difference between AH+ and HC was not as significant (t = −1.558, p = .124).

**Figure 4.**
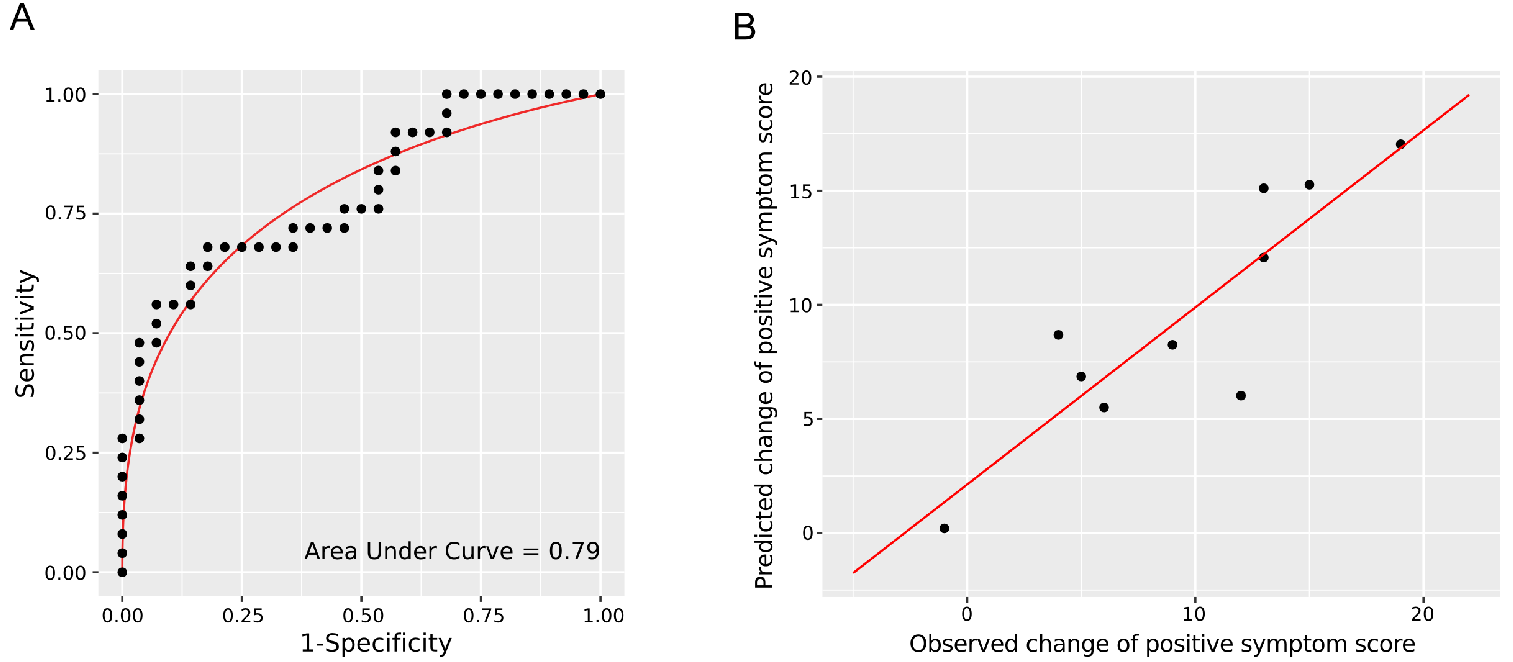
A. Receptive Operation Curve (ROC) of the logistic regression model to classify AH+ from general schizophrenic patients. The black dots represent the sensitivity and 1-specificity pairs obtained by applying different threshold of probability. The red curve shows a smoothed ROC curve. The Area Under Curve is 0.79; B. Correlation between the observed and predicted improvements of positive symptom scores in AH+ patients (R^2^ = 0.77). The observed changes reflects positive score difference before and after 2-months’ medication, and the predicted changes are from a linear regression model with PMN-hippocampus functional connectivity strength, DUP, and age as regressors.

Longitudinal analysis did not show significant group by time interaction on the precuneus-hippocampus connectivity (F = 1.226, p = .303). However, the changes of precuneus ROI-hippocampus connectivity exhibited negative correlation with the reduction of positive symptoms of BPRS (r = −.577, p = .081) in the AH+ group. In AH− group, there was no significant correlation between clinical symptom changes and alternations of the precuneus-hippocampus connectivity.

### Predict AH symptom using functional connectivity strength

To model the contributions of FCS features to predict AH+ in schizophrenia, we conducted a stepwise logistic regression between AH+ and AH− group. We included mean weight of whole PMN, mean weight of precuneus ROI (indicating within network FCS), precuneus-hippocampus connectivity, and other covariates such as DUP, education level, age, gender in logistic regression with baseline data. Results demonstrated that only the mean weight of precuneus ROI (beta = −.071, p = .010) and PCC-Hip connectivity (beta = −4.249, p = .019) survived in the model (Supplementary Table 2).

In the ROC analysis of the simplified model (mean weight of precuneus ROI and precuneus-hippocampus connectivity) to distinguish AH+ from all schizophrenic patients (Figure 4A), the AUC of the simplified model was 0.793. The leave-one-out cross-validation test estimated the prediction accuracy of the model as 0.731, with a sensitivity of 0.750 and a specificity of 0.708.

We further built a model to investigate whether the functional connectivity features at the baseline scan help to predict improvement of clinical symptoms in AH+ patients. The mean weight of the precuneus ROI, the precuneus-hippocampus connectivity, and demographical variables were used as regressors to predict the change of positive scores of BPRS of AH+ patients after treatment. The precuneus-hippocampus connectivity, DUP, and age survived the stepwise regression, and explained 77% of the variance of positive score changes (F = 6.936, p = 0.022, Figure 4B). A leave-one-out cross-validation yielded an R^2^ of 0.433 (F = 6.115, p = 0.039). As a contrast, the same model selection and cross-validation procedure failed to yield a significant predictive model for AH− patients.

## Discussion

With a unique sample of first-episode drug-naïve schizophrenia patients, this study identified AH+ specific dysfunctions in the PMN, supporting our hypothesis regarding PMN deficits in AH. Moreover, precuneus (a core region in the PMN) and its functional connectivity with hippocampus showed significant contributions in identifying AH+ from general schizophrenic patients and in predicting clinical improvements of AH+ patients after treatment, suggesting novel and precise clinical predictors.

### PMN’s deficits distinguish AH+ from general schizophrenic patients

Our findings help to differentiate neural network deficits underlying AH symptoms from those underlying general schizophrenic symptoms. First, AH+ patients showed the weakest FCS within the PMN, with the peak difference appeared at the precuneus. The FCS of this region then increased with the alleviation of AH symptoms after a 2-month medication. In the HC and AH− groups, however, this region showed no significant change between the baseline and follow-up scans. Second, the functional connectivity of the precuneus ROI was negatively correlated with the severity of AH symptoms. Third, our regression models have demonstrated the predictive power of the PMN-related functional connectivity features for identification and clinical prognosis of AH+ patients. These results suggest that the PMN deficit is a neural network marker for AH symptoms.

### A novel view of DMN’s role in AH

This study further distinguishes the roles of the PMN from the DMN in schizophrenia. An intrinsic connectivity network centered at the precuneus has been separated from DMN by multiple methodologies (Doucet *et al.*, 2011; Power *et al.*, 2011; Yeo *et al.*, 2011), and it has been recently linked to familiarity labelling by Gilmore’s work (Gilmore *et al.*, 2015) and therefore named as PMN. However, the role of the PMN in mental disorders has not been clarified. Previous neuroimaging studies have sometimes accounted for alternations of the precuneus to the abnormality of the DMN or other networks (Sheline *et al.*, 2010; Manoliu *et al.*, 2014; van Lutterveld *et al.*, 2014), although anatomical and functional studies have indicated the separation of precuneus from the DMN (Buckner *et al.*, 2008; Zhang and Li, 2012), and a few studies have implied the relevance of the precuneus to AH (Sadaghiani *et al.*, 2009; Nenadic *et al.*, 2010; Wolf *et al.*, 2011; Allen *et al.*, 2012). Our study further clarified the functional boarder between PMN and DMN by demonstrating a contrast between the PMN and DMN regarding AH symptom in schizophrenic patients.

### Possible role of PMN underlying AH

The results on the interaction pattern between the precuneus ROI and hippocampus further explained the role of the PMN in AH. The decreased FCS between the right precuneus and bilateral hippocampi reflects how deficits in the PMN affect memory processing in AH. Studies in schizophrenia patients have suggested that dysconnectivity of large-scale networks may result from aberrant modulation of N-methyl-D-aspartate (NMDA)-dependent plasticity (Friston and Frith, 1995; Stephan *et al.*, 2006; Stephan *et al.*, 2009). Abnormal antagonist of NMDA receptor, such as ketamine, may cause schizophrenic-like symptoms including perceptual aberration. The dysfunction between PMN core region and hippocampus in present study support the possible involvement of memory system in AH. Unstable signal fluctuations in parahippocampal (Diederen *et al.*, 2010) and dysconnectivity patterns of hippocampal complex (Amad *et al.*, 2014; Lefebvre *et al.*, 2016) have been evidenced during hallucination state, indicating a failure of cognitive suppression of irrelevant memory into consciousness. The PMN, which involve in the encoding and retrieval of memory, may correspond to the ability of inhibiting irrelevant memory and reinforcing novel-materials learning in real context (Gilmore *et al.*, 2015). This interpretation is supported by a recent TMS study reporting that improvement of associative memory performance was achieved by enhancing connectivity between the left hippocampus and the precuneus/retrosplenial cortex (Wang *et al.*, 2014). Another study also showed that semantic memory was associated with increased connectivity between the cuneus/precuneus and left hippocampus (Sormaz *et al.*, 2017).

Therefore, we propose the role of PMN in generation of AH as a cognitive supervisor, further refining the “unstable memory” hypothesis. The deficit of PMN may produce erroneous activity to guide AH+ patients to pay more attention to familiar features of exogenous objects and retrieval of autobiographical memory that is irrelevant to the circumstance in reality. With the unstable intrinsic fluctuation of the PMN, the sudden appearance of AH would become inevitable. This explanation extends existing neural network theories of AH (Northoff and Qin, 2011; Jardri *et al.*, 2013) by connecting memory-related dysfunction to its underlying neural network, and hence provides more specific targets for potential treatments.

### Limitations

A limitation of the study is that the state of the participants during scanning was not recorded or controlled so that AH may occur, which did not allow us to identify the state-related biomarkers. A paradigm that better constrains mental states and a post hoc questionnaire should be used in further studies to overcome this issue. Another limitation is the relatively high missing follow-up rate due to high population mobility in the city. High missing follow-up rate may partially limit our statistical power in the follow-up analysis, and this factor should be taking into account when interpreting the longitudinal findings, especially the negative ones.

## Conclusions

This study proposed and examined a novel hypothesis that deficits in intrinsic activity of the PMN and its interaction with hippocampus are specific to AH symptoms. While not rejecting the contribution from dysfunction in the DMN, this study helped to explain how dysfunction in the memory system cause AH and further identified a novel neuroimaging marker and treatment target for AH. Future works should focus on delineating the detailed deficit in the PMN and confirming treatment effects via intervention on PMN activity.

## Acknowledgments

The authors have no conflict of interest to declare.

